# Single-cell analysis of *Plasmodium falciparum* transcripts after drug perturbation identifies feedback regulation as well as increased transmission potential

**DOI:** 10.64898/2026.05.27.728291

**Authors:** Karla P. Godinez-Macias, Jaeson Calla, Kristen Jepsen, Elizabeth A. Winzeler

## Abstract

Gene expression analysis in malaria parasites has been used to define transcriptional regulatory networks but has been used less frequently to characterize parasite response to drug treatment or to show how parasites may evade killing. Here, we applied single-cell RNA sequencing (scRNA-seq) to hundreds of thousands of individually infected asynchronous red blood cells to evaluate the parasite’s response to treatment with three chemotypes that can be used for treatment (artemisinin) or prophylaxis and treatment (atovaquone, ganaplacide). We found that each treatment gave rise to different cell populations with different transcriptional profiles. Comparing single cell transcription patterns in compound-treated cells, to transcript patterns observed previously with synchronized cells showed an enrichment of cells expressing gametocyte-associated genes after artemisinin treatment but fewer lifecycle perturbations after treatment with the two other compounds. In contrast, bulk analysis showed an enrichment of pyrimidine biosynthesis transcripts for atovaquone treatment. Our results show that scRNA-seq may be used to profile diverse drug responses across many lifecycle stages and to potentially classify drug classes.

**Importance:** Determining the mechanism of action (MOA) of compounds with antimalarial activity remains a key activity in both drug development and drug resistance studies but remains challenging for some chemotypes. Here we highlight the potential of single cell transcriptional sequencing to augment the process of MOA deconvolution. We develop a new analytical pipeline that involves comparing single cell transcription patterns to existing profiles from synchronized parasites to comprehensively characterize life cycle stage enrichments that may be observed after chemical perturbations. We also show that transcriptional feedback regulation may be present for some drug classes.

## Introduction

Phenotypic screening has led to the discovery of tens of thousands of novel antimalarial chemotypes, many of which could map to a new validated target. In many cases, *in vitro* evolution and whole genome analysis of the resulting strain has been used to map compounds to targets but it can be time-consuming and, in some cases, resistant parasites cannot be obtained or resistance data cannot be interpreted(1). Gene expression profiling has been used to elucidate transcriptional regulatory networks and to functionally assign uncharacterized genes to networks in both malaria parasites(2–5) and in other species(6–8). In theory, transcriptional profiling could be used to determine a target or mechanism of action (MOA) of uncharacterized small molecules, especially in cases where a metabolic block leads the parasite to compensate by transcriptionally upregulating the genes in the blocked pathway (*e.g.*, upregulation of the folate biosynthetic pathway after treatment with an antifolate)(9, 10). This principle of comprehensively relating bioactive compounds to their target pathways and proteins has been demonstrated using *Saccharomyces cerevisiae* by clustering groups of transcriptional responses with similar expression profiles(10). In addition, even if a specific pathway is not revealed, transcriptional profiles could be used to categorize drugs with the same MoA(10) (*e.g.*, all antifolates should give similar antifolate profiles), or uncover underlying mechanisms of phenotypes such as drug resistance(11).

Some of the challenges with applying transcriptional profiling for compound MOA studies in malaria parasites include synchronizing cells and determining the optimum cell type to conduct expression profiling(12). Furthermore, determining the time and concentration of drug exposure, in the case of treatment response, adds an extra layer of liability. Hypothetically, applying single-cell transcriptional analysis could help overcome this problem by allowing profile analysis in a variety of different cell types all at once(12). Single cell RNA sequencing (scRNA-seq), including the Chromium platform from 10× Genomics™, allows a continuum of cell types to be analyzed simultaneously. In this method, single parasites from different stages are individually encapsulated in a GEM droplet(13) and are subsequently barcoded and their transcriptomes sequenced and deconvoluted.

Here we assessed the use of 10× Genomics™ scRNA-seq Chromium platform for drug target/MOA identification. We performed transcriptomic profiling on human red blood cells infected with *P. falciparum* asexual blood stage (ABS) parasites, treated with one of artemisinin, atovaquone, and GNF179, a close analog of ganaplacide (also known as KAF156). Atovaquone is a naphthoquinone scaffold targeting *Plasmodium* cytochrome bc-1 complex(14) as part of the respiratory chain complex. GNF179 is a very close analog of ganaplacide, and both appear to work through disrupting protein trafficking mediated by *Pf*CARL, an ER-localized transmembrane protein(15–17). Lastly, while no definitive target is known for artemisinin (sesquiterpene lactone class)(18), it activates free radicals in the presence of heme leading to parasites’ death(11, 18).

Next, we developed a customized bioinformatic pipeline that not only identifies classes of transcriptionally regulated genes when comparing treated versus non-treated parasites, but also distinguishes changes in numbers of different cell types across the parasites lifecycle by searching for cell type dependent transcriptional signatures that were previously discovered using gene expression analysis of synchronous parasites from different lifecycle stages. When comparing the transcriptional response distinguishing treated and non-treated parasites, we observed disruption of parasite development after drug exposure as well as increased commitment to gametocytogenesis. Our atovaquone data also suggests that specific upregulation of the inhibited pathways may be observed in malaria parasites.

## Results

### Profiling drug treated Plasmodium falciparum asexual blood stage parasites

To perform cell profiling, three independent biological replicates of asynchronous asexual blood stage *Plasmodium falciparum* Dd2 parasites at 4% parasitemia were added to four different flasks per replicate; one flask was maintained as an untreated control, and the remaining flasks were treated with one of three antimalarials as described in **Figure 1a**. Next, 100nM of artemisinin, 50nM of atovaquone, and 100nM of GNF179 were added to one of the three flasks, respectively. To avoid sequencing mostly uninfected cells, after 48 hours of drug exposure, infected cells from both the drug-treated and control samples were extracted using magnetic-activated cell sorting (MACS) columns separation (replicate 2 and 3). Samples were then prepared for single cell capture, barcoding, library preparation, and transcript sequencing using 10× Genomics™ technology (see **Methods** for details). Following sequencing, reads were mapped to a custom reference genome of human (GRch38.p13) and *P. falciparum* Dd2 (v62), with on average 2,333 reads per cell (**Table 1**).

**Figure 1.**
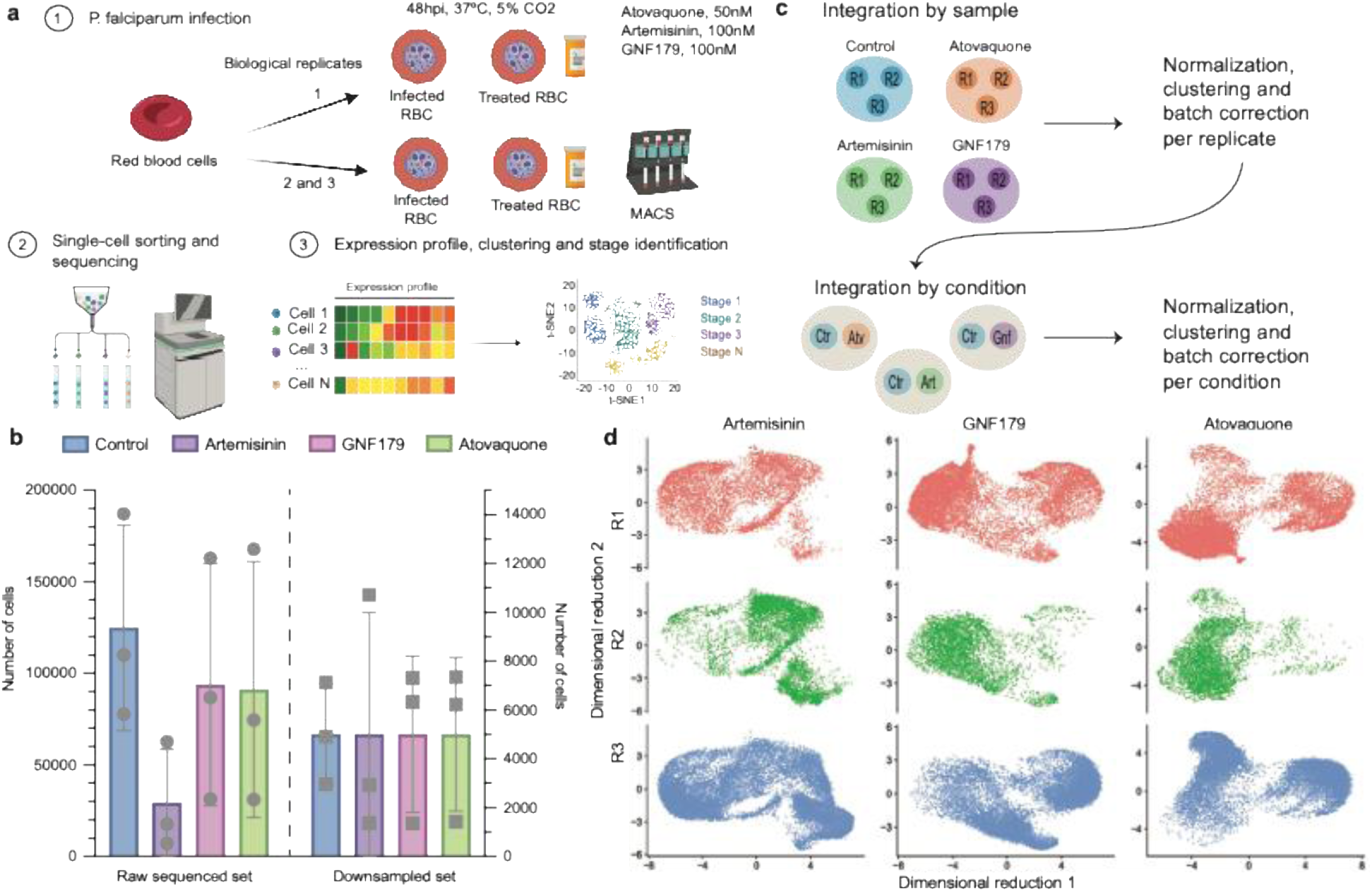
Experimental design and processing of *P. falciparum* asexual parasites. a) Experimental design workflow. Steps followed to study drug treatment in *Plasmodium falciparum* Dd2 parasites. Three biological replicates were performed; human red blood cells were infected with Dd2 parasites, one was left untreated (control) and three separated iRBC were treated with atovaquone, artemisinin and GNF179 at listed IC_50_ concentrations. After 48 hours post infection, parasites were prepared for 10× Chromium single-cell protocol and sequenced using NovaSeq S4 flow cell (NovaSeq 6000 system). A custom bioinformatic workflow was followed, using a combined human and *Plasmodium* genome for alignment and transcript quantification. Downstream analyses were performed on *Plasmodium* cells only. **b) Number of raw cells and downsampled cells across samples.** Histogram contrasting the total number of parasites sequenced across control (blue), artemisinin-(purple), GNF179-(pink) and atovaquone-treated (green). **c) Integration and correction workflow.** Pre-processing steps followed to integrate control cells and drug-treated cells. The package tool Seurat(21) was used for the processing and analysis of cells. Replicates for each sample were integrated as a single object (control R1-3, atovaquone R1-3, artemisinin R1-3 and GNF179 R1-3) using the merge option, respectively. Objects were subject to a normal single-cell pipeline correcting using Harmony by replicate (normalization, cell cluster obtention, batch correction). The corrected objects, were integrated using Seurat’s merge function, having *x* as the control and *y* as the drug-treated (artemisinin, atovaquone, or GNF179), and subject to a standard single-cell pipeline and batch correcting according to the infected or treated condition. Genes in hypervariable regions of the genome and outlier cells were filtered. **d) Cell integration and reproducibility.** Dimensional reduction map (UMAP) for control and drug-treated integrated cells, as determined by Seurat(21) DimPlot. UMAP showing cells split by replicate (1, orange; 2, green; and 3, blue) across cell-treated cells (artemisinin, GNF179, and atovaquone) were plotted.

**Table 1.**
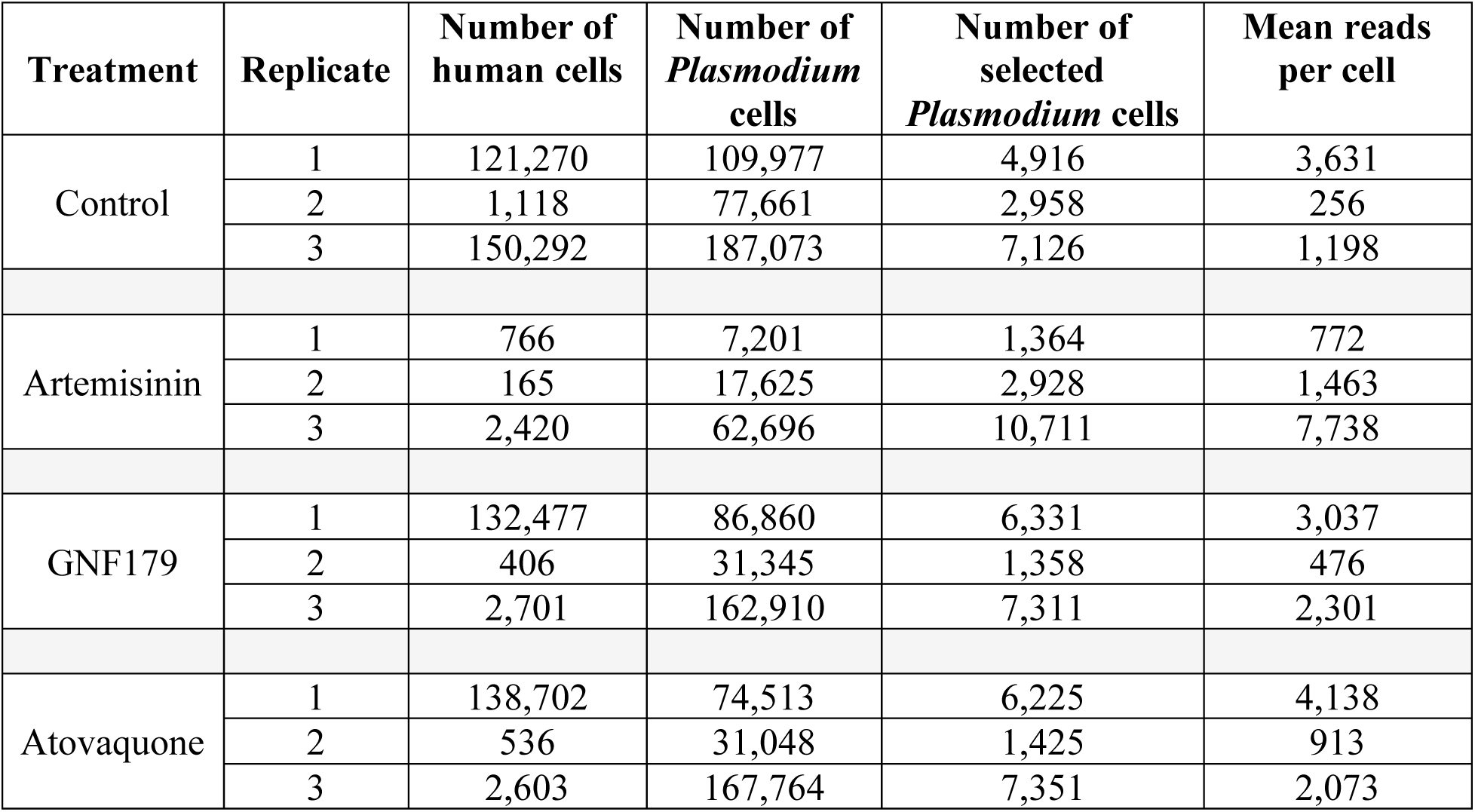
Estimated number of cells and total number of cells selected for this study. Estimated number of cells that were obtained from human and malaria parasite genomes as calculated with CellRanger. Total number of cells used for analyses after random selection for each sample are shown.

Considering the use of magnetic columns in two replicates, we were expecting to see an enrichment in sequenced malaria parasite cells compared to human cells. However, according to our data, this did not always hold (**Table 1**). For example, transcripts in the third replicate control were nearly evenly distributed between human and malaria cells, and drug-treated samples showed higher human cells. The presence of human transcripts with the same barcode as parasites is expected given that parasites are within human cells and although red blood cells (which lack nuclei) may lose transcripts as they mature, some retention is expected. Additionally, we observed large differences in the total number of malaria cells recovered per sample and across replicates (**Fig. 1b**). For example, 4.28× fewer cells were obtained after artemisinin treatment, compared to the control. This likely reflects differences from drug treatment. On average, we obtained 124,903.67 *Plasmodium* cells for controls (non-treated), 29,174 cells for artemisinin-treated cells, 91,108.33 cells for atovaquone-treated cells, and 93,705 cells for GNF179-treated cells. To avoid skewing the results with oversampled groups and to ensure gene expression comparisons across conditions, we randomly selected a total of 15,000 cells per condition (**Table S1**, **Fig. 1b**). Except for artemisinin-treated cells, the downsampled set showed lower variability (average SD 3,357.82) compared to the full set (average SD 55,402.29), suggesting fewer biases towards a particular condition or replicate.

To evaluate the data, we first integrated total parasite Counts per 10 Thousand log-transformed (as represented by CP10K) per protein coding genes across all cells and compared these aggregate results to rankings previously obtained with a historical mRNA abundance study of different parasite synchronous lifecycle stages(3). The sets were similar with 4,440/5,211 transcripts detected above background (≥50 CP10K) in the scRNA-seq control cells and 4,093/4,672 genes above background in the historical dataset (20 units) considering only ABS samples. Background expressed genes in scRNA-seq average control included genes involved in antigenic variation (GO:0042783, Bonferroni *P* = 2.90×10^-8^) as well as meiosis, as expected, and included examples such as CSP. Likewise, genes involved in gliding motility and axoneme function were excluded from the detected ABS transcripts in the historical dataset. This comparison largely showed that our scRNA-seq results were as expected with 65% (*n* = 23) of the most abundant transcripts in our control cells (>10,000, *n* = 35) among the most abundant transcripts in the other study (panova < 5×10^-5^, nprobes ≥ 5) (**Table S2**). For example, we found the exported protein 1 (PF3D7_1121600), HSP70 (PF3D7_0818900), and early transcribed membrane proteins ETRAMP2 (PF3D7_0202500) and ETRAMP11.2 (PF3D7_1102800) were among the top 1% most abundant transcripts confidently detected (*p* < 0.05) in asexual blood stage samples from both studies. We observed a few cases with large discrepancies in abundance and these may be due to strain-specific differences. For example, the most notable was *Pf*KAHRP (PF3D7_0202000, knob-associated histidine-rich protein), which was barely detected in the newer single-cell study. Dd2 parasites are known to carry a structural deletion of the KAHRP gene(19), which provides a reason for the marked difference. Strain-specific differences also likely accounted for discrepant abundance rankings for other antigens, such as S-antigen (PF3D7_1434500), which was highly expressed only in the scRNA-seq samples. This is likely due to low protein sequence identity between Dd2 and 3D7 for the S-antigen(20). For most conserved housekeeping genes that were adequately probed in both datasets, correlation was excellent (*r =* 0.61, Spearman). Examples include actin I (PF3D7_1246200), myosin A (PF3D7_1342600) and ribosomal protein S8 (PF3D7_0707900).

Confident with mapping quality with our scRNA-seq dataset, we sought to investigate the transcriptional changes after drug treatment at the single cell level. As described in **Figure 1c**, we combined the single-cell sets from the three replicates per condition and performed a standard single-cell analysis (normalization, feature selection, scaling, dimensional reduction), followed by combining cells from the control and one of the drug-treated samples (artemisinin, atovaquone, or GNF179), using Seurat(21) and Harmony batch correction on the replicate number and treatment. We also filtered out transcripts mapping to noncoding genes and hypervariable gene families to prevent mapping artifacts(22). In general, we observed that cells were integrated as expected across replicates (**Fig. 1d)**. In most cases, the third replicate showed the highest number of unique cells for all treatments, with an average of 1,192.08 cells, compared to the first and second replicates (**Table S3**). This was expected given the third replicate had the highest estimated number of recovered cells. Nevertheless, we observed cells from all replicates present across the combined single-cell sets, confirming the overall effectiveness of the integration.

### Plasmodium cells showing lifecycle markers are different after drug treatment

There are two competing hypotheses about what happens after parasite drug treatment which could be resolved by single-cell analysis. The first is that there is a shift in the distribution of established cell types. The second is that new cell types with a different gene expression profile emerge. To test these two hypotheses, we first performed cell type clustering analysis using single-cell resolution expression data with the assumption that each cluster should be representative of a known intraerythrocytic parasite developmental cycle (**Fig. S1**). We applied the “shared nearest neighbor” algorithm (SSN), which is based on grouping cells that share the most cells with equal gene expression profile in a network graph and separating those cells having few or no neighbors(23). We hypothesized that we would be able to detect clear substages (*i.e.*, ring, trophozoite, schizont, merozoite), as well as parasites undergoing the developmental processes. From this clustering, we obtained a total of 13 substages (clusters) for parasites treated with artemisinin with an average of 2,255 cells per cluster, atovaquone-treated parasites were distributed across 18 substages with an average of 1,666 cells per cluster, and 24 substages were found after GNF179 treatment having an average of 1,250 cells per cluster (**Table S3**).

Upon closer inspection of the ABS substages, we found some cases with low number of cells per stage or with noticeable cell differences between replicates per stage. This included substages 14–18 and 14–24 for atovaquone and artemisinin-treated cells, respectively. The difference in total number of clusters could be related to the parasite clearance effects. For example, artemisinin’s fast killing profile should prevent early parasites from developing, but those in later stages may have undergone the full cycle. For GNF179 profile, on the other hand, we expected less stages throughout the life cycle because of its early erythrocytic and gametocytocidal activity(15). The clustering cell differences could also be due to an artifact during the integration steps(24), and were thus filtered out due to a lack of statistical confidence. This resulted in thirteen, ten, and eight ABS substages for artemisinin, atovaquone, and GNF179 treated parasites, respectively.

We next aimed to identify which asexual developmental stages were present in the cell clusters (**Table S4**). To accomplish this, we used Seurat’s *FindMarkers* function to identify *P. falciparum* genes with the highest expression in the cluster of interest compared to the rest. To tie our Seurat single-cell clusters to specific parasite cell types, we matched Seurat marker genes to genes that have previously been shown to be upregulated in specific purified, synchronous cell types (e.g. rhoptry genes have been shown to be upregulated in schizonts(3)). For a detailed description on cluster assignment, please refer to **Methods**.

This matching showed a high level of overlap. For example, cells in cluster 3 for GNF179 tended to show strongest expression for the same sets of genes that are highly expressed in purified “late schizont” parasites described previously (*P* = 6.65×10^-12^) according to the hypergeometric test. Here Seurat single-cell marker genes included many genes encoding merozoite surface proteins and rhoptry neck proteins which have previously been tied to the late schizont phase. Using this matching process we assigned single-cell clusters to ring, trophozoite, schizont, merozoite, and gametocyte types (**Fig. 2a**).

**Figure 2.**
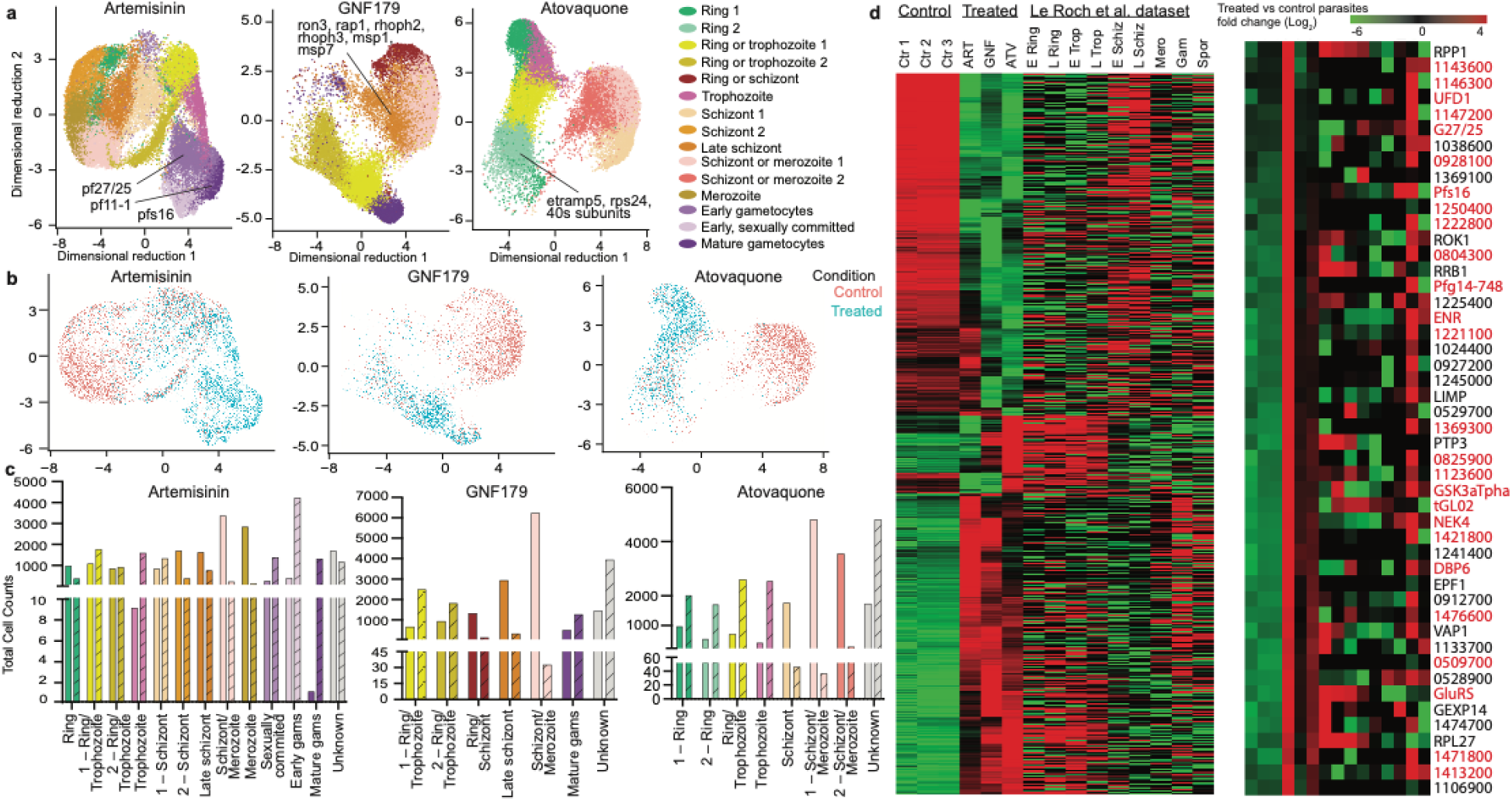
Transcriptional profiling of asexual *Plasmodium* parasites relative to after drug treatment. a) Parasite cluster stages after artemisinin, GNF179 and atovaquone treatment. Dimensional reduction of high-quality cells relative to after drug treatment were visualized using Seurat’s DimPlot function. Identified clusters (see **Methods**) were color-coded and displayed across treatments to facilitate comparisons. Unidentified (N/A) clusters were filtered for clarity. **b) Cluster membership for treated vs. control cells.** Dimensional reduction UMAP plot for each treatment is shown. Drug-treated parasite cells are colored in teal and control parasites are shown in salmon. Plot was generated using Seurat DimPlot and manually corroborating with read counts. Filtered clusters were removed for simplicity **c) Number of cells recovered per ABS cluster stage after artemisinin, GNF179, and atovaquone treatment.** BarPlot showing total number of cells recovered in the single-cell experiment after each treatment. The *X-*axis shows the cluster substages recovered per treatment, and the *Y-*axis the total number of cells per predicted substage. Bars are colored by the corresponding stage as in Fig. 2a. Recovered cells in the control are shown with fully colored bars while drug-treated cell counts are colored with diagonal lines. d) Hierarchical clustering of single-cell transcript counts and mRNA abundance of *Plasmodium* lifecycle. Clustering performed by gene expression patterns on total raw transcript counts (UMI in Seurat) for drug-treated versus control parasites in the single-cell study. The total number of transcripts mapping in the Le Roch study(3) were mapped for the complete lifecycle and ordered based on the single cell clustering. A total of 1,635 significant genes (*P* = 5.10x^-5^), highly expressed (>100 transcript counts), and confidently mapped between strains are shown. Color scale indicates log_2_ fold-change from-6 to 4. Erythrocytic substages are labeled as follows: Trop represent trophozoites, Schiz represents schizonts, and Mero represents merozoites. Gametocytes are indicated with the label Gam and sporozoites with the label Spor. In addition, parasite substages at early development phases are represented by an “E” and fully developed are represented by an “L”, prior to the corresponding substage (e.g. E trop are parasites at early trophozoite stage). A zoom in on gametocyte/sporozoite gene enrichment with artemisinin treatment in the single-cell study is highlighted in dark grey and displayed. Genes highlighted in red, with the symbol or 3D7 id, indicate genes represented in gametocyte or sporozoite according to Le Roch study.

Next, we compared the distribution of predicted cell types (*i.e.*, ring, trophozoite, schizont, merozoite, and gametocytes) before and after drug treatment (**Fig. 2b**). As expected, not every substage was present after all treatments. For example, no gametocytes were observed after atovaquone treatment, and no rings were captured after GNF179 treatment. For both GNF179 and atovaquone, drug treatment resulted in a general loss of predicted cell types. For artemisinin however, we observed the emergence of predicted gametocyte stages which were not as prevalent before treatment. This is consistent with artemisinin being less active against parasites that are committed to sexual development (25).

To quantify this we estimated the numbers of different cell types after normalization before and after drug treatment using their predicted assignments (**Table S6, Fig. 2c**). From this, artemisinin showed a significant reduction in cells with predicted schizont/merozoite profiles (3,293 unique cells recovered in the control single-cell set to 152 unique cells recovered in drug-treated parasites) and merozoite profiles (2,762 to 51), but a significant increase in cells with trophozoites (9 to 1,511) and gametocyte profiles (316 to 4,157 for predicted early gametocytes, 175 to 1,236 sexually committed, and 1 to 1,236 in mature gametocytes types). In contrast to artemisinin, GNF179 showed a major decrease primarily in predicted late schizonts (2,933 to 323) and schizont/merozoite parasites (6,225 to 36), consistent with its inhibitory profile(26). Atovaquone showed a decrease in mature schizont parasites (1,733 cells to 53 cells) and schizont/merozoite types (4,820 to 42 cluster 1, and 3,547 to 113 cells cluster 2), similarly to its well-documented slow acting profile(27). Notably, after atovaquone treatment there was a ∼3× fold increase of predicted “unknown” cell types going from 1,706 to 4,811 recovered cells across clusters five, seven, and ten.

### Gametocyte-specific gene expression is induced after artemisinin treatment

We next sought to corroborate the single-cell analysis using an alternative approach. Here we summed CP10K transcripts per single-cell across conditions for all sequenced single cells and all genes and then performed differential gene expression (DEG) analysis (**Table S7**) on the resulting set. We obtained a total of 1,856 statistically differentially expressed genes (up-and down-regulated*, P-adj* ≤ 5×10^-5^) after artemisinin exposure compared to non-treated parasites, 1,427 genes, and 1,670 genes changing after GNF179 and atovaquone exposure, respectively. We next asked if genes that were upregulated after artemisinin treatment in the single-cell analysis would also show upregulation in previously published gametocyte data. We selected genes with more than 50 transcript read counts (UMI in Seurat) in at least one single-cell condition (control or drug), a log_2_ fold-change of at least ±1, and a confident mapping between the 3D7 and Dd2 strains. This resulted in a total of 1,635 unique genes. We then plotted their drug-related gene expression profiles alongside the corresponding expression level for previously published synchronous malaria parasite lifecycle asexual, gametocyte and sporozoite stages(3) (**Fig. 2d**). From this, it could be seen that artemisinin induced a strong upregulation of genes upregulated in purified gametocytes, such as *g27/25*, *pfs16* and *pf11-1* (*P* = 1.90×10^-89^), and in sporozoite parasites (*e.g., csp*, *pdi-8*, *trap*) (*p* = 5.74×10^-5^) using as baseline the Le Roch et al. gene expression set.

### Gene ontology enrichment shows induction of pyrimidine biosynthesis pathway after atovaquone treatment

The analysis above did not address the alternative hypothesis that drug treatment could create new classes of cell types that are not observed without drug treatment. In fact, we did observe some drug-induced cell clusters which were not present before treatment and that did not show clear enrichments of known cell parasite lifecycle markers. Thus, here, we used an alternative approach. After integrating the number of transcript reads per gene for all single-cells, we took the list of differentially expressed genes resulting in 409, 188, and 462 unique upregulated genes in atovaquone, GNF179 and artemisinin, respectively (log_2_ fold-change > 1) and 506, 571 and 550 downregulated genes (log_2_ fold-change <-1). These groups were then subjected to gene enrichment analysis. From this list, we found majority of upregulated genes were exclusive to artemisinin (*n* = 273) or atovaquone (*n* = 229) treated parasites (**Fig. 3a**). This analysis confirmed what was observed above: for genes upregulated only in artemisinin, we found gametocyte and sporozoite markers including for gamete egress proteins (*pfgep*, *pfgep1)* and sporozoite invasion-associated proteins (*pfcsp* PF3D7_0304600, *pfsiap-*2 PF3D7_0830300). Enrichment analysis showed signatures of gamete formation, mobility and transmission with artemisinin (*P* = 1.34–4.80×10^-2^, Bonferroni corrected). Some examples include cell gliding (5 out of 14, GO:0071976), cilium movement (4 out of 7, GO:0003341), and crystalloid formation (6 out of 18, GO:0044312). This was further confirmed when comparing the membership of similar gene functions in the Le Roch et al. clustering set, showing the highest peak in gametocyte and sporozoite genes was with artemisinin (**Fig. 3b**). In addition, enrichment analysis showed a strong translation and ribosomal subunits (*P* = 9.22×10^-21^ to 4.34×10^-2^) for genes downregulated after artemisinin treatment.

**Figure 3.**
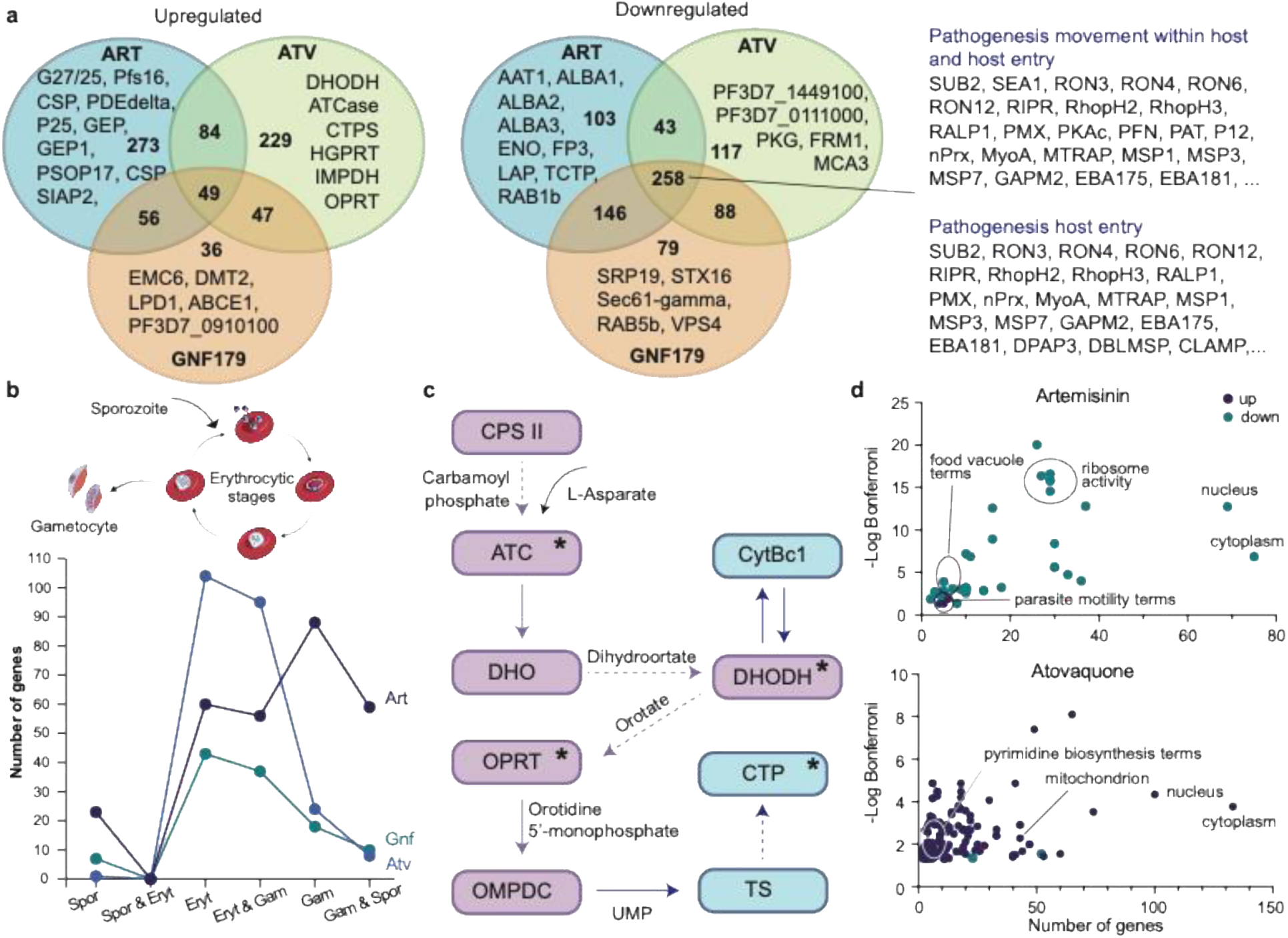
Differential expression enrichment. a) Venn Diagram with differentially expressed genes log_2_ fold of at least ±1. Significant genes from **Table S4** were compared across treatments. Upregulated genes for artemisinin (*n* = 462), GNF179 (*n* = 188) and atovaquone treated parasites (*n* = 409) are shown according to the treatment overlap. Downregulated genes for artemisinin (*n* = 550), GNF179 (*n* = 571) and atovaquone (*n* = 609) are shown. Genes related to the mechanisms of action or suspected function are labeled. **b) Pyrimidine biosynthesis pathway with significant genes highlighted.** A simplified version of the pyrimidine biosynthesis pathway(43) is shown with related genes colored in purple (CPS-II, ATC, DHO, DHODH, OPRT, OMPDC). Additional interaction (TS, CTP) or dependent enzymes (CytBc1) are highlighted in blue. Dashed arrows indicate the presence of a gene in the pathway that was significantly upregulated in the single cell study, while double-arrows indicate a direct relationship between a gene in the pathway (DHODH) and an external gene (CytBc1)(43). For representation, a star is included in the enzyme name that was mentioned in the text. **c) Total of significant genes expressed mostly in the malaria erythrocytic, gametocyte and sporozoite stages.** A schematic of the malaria parasite stages considered are shown. A total of 243 significant genes for atovaquone, 288 for artemisinin and 120 for GNF179 were among the Le Roch developmental classification set. A histogram line graph comparing the total genes matching to each stage, or between stages, across treatments is shown. Atovaquone is labeled in royal blue, artemisinin in dark blue, and GNF179 in green. **d) GO enrichment**. Scatterplot showing for artemisinin (top graph) and atovaquone (bottom graph). GNF179 did not show statistically significant terms and thus is not included. The terms for biological processes, molecular functions, and cellular components are shown together, with enriched terms from the upregulated set marked in dark blue and the downregulated set marked in green. Total number of genes appearing with each term is shown in the *X* axis and the-Log Bonferroni *P*-value is shown in the *Y* axis. Terms with significant *p-*value or relevant mechanism pathway are labeled in each plot.

Interestingly, among the upregulated genes present only after atovaquone treatment we observed three of the six enzymes involved in the *de novo* pyrimidine biosynthesis(28). These were ATCase (PF3D7_1344800), DHODH (PF3D7_0603300) and OPRT (orotate phosphoribosyltransferase, PF3D7_0512700) (**Fig. 3c**). Another upregulated enzyme, CTP synthase (PF3D7_1410200), while not part of the same essential pathway, is a direct product of the same pathway(28). These data clearly relate to the downstream effect of atovaquone causing disruptions in the pyrimidine biosynthesis pathway. Unbiased GO analysis (**Table S8**) showed enrichments in the pyrimidine biosynthesis pathway and required reactions (*P* = 2.08×10^-3^ to 5.58×10^-5^), in agreement with the parasites mechanism of response for this drug(27). These data indicate that transcriptional feedback regulation is occurring for atovaquone (**Fig. 3d**).

Unfortunately, GNF179 did not show any significant enrichment in the sets of up and down regulated genes, which could be due to the lack of specificity in the GO comparisons rather than a true lack of signature. The lack of significance metabolic enrichment with the imidazopiperazine class has been observed with other methods as well(29). These compounds are known to act against gametocytes as well which could be why fewer gametocyte signatures were observed than with artemisinin. We did see seven genes (TryThrA, SRP19, SMS1, SHLP1, Sec61-gamma, PDI-14, ACS3) associated with the endoplasmic reticulum (ER) some of which are involved in protein trafficking to or from the ER, including SRP19 (PF3D7_0210000), RAB5b (PF3D7_1310600), Sec61-gamma (PF3D7_1216300) and STX16 (PF3D7_1243000), although the enrichment was not significant.

## Discussion

The question of whether malaria parasites respond transcriptionally in a way that can be used to glean compound mechanism of action has a long history of debate in the malaria community. When the *P. falciparum* genome was first sequenced, fewer transcription factors were identified than in other species, suggesting a paucity of transcriptional control mechanisms(30, 31). Early gene expression studies with microarrays promoted the idea that gene expression was cyclical in nature with each gene being turned on a specific, “just-in-time” format in the asexual blood stage(2). Later, more transcription factors were identified suggesting a more nuanced view(32). More recently, Luth et al.(33), identified many mutations in transcription factors in their studies on the genetic basis of drug resistance. On the other hand, studies looking for transcriptional feedback regulation after drug treatment in malaria parasites have seldom yielded solid results. Studies looking at gene expression with 31 different small molecules did not yield clear mechanism of action(34). Early studies with antifolates were also unclear about whether specific feedback regulation exist(35).

Evidence does suggest that transcriptional feedback regulation should exist. Previous studies have demonstrated that metabolic profiling can reveal the drug’s mechanism of action by looking for changes in metabolic pathway intermediates after drug exposure(36, 37). These studies were able to show specific, reproducible upregulation of pyrimidine pathway intermediary metabolites after atovaquone treatment(36) as well as upregulation of purine metabolites after antifolate treatment. This was also confirmed by Simwela et al.(29). Our data suggest that the changes in pyrimidine pathway enzymes that are seen in the metabolome in malaria parasites derive from changes in transcripts.

The role of specific transcription factors in modulating these changes remains to be explored. It is likely there may be some role for AP2-G (PF3D7_1222600), the master gametocyte transcription factor(38), in modulating the response to artemisinin and other drugs that cause cellular stress. AP2-G is one of the most frequently mutated genes in long term drug selections, and while this may be related to the loss of gametocyte-production abilities in parasites kept in long term culture, we have noticed that it is frequently co-mutated with the master drug regulator, *Pf*MDR1(33). Nevertheless, upregulation of AP2-G was modest (∼2×), although we observed strong upregulation of two poorly characterized transcription factors that are essential for transmission stages, AP2-O3 (PF3D7_1429200) and AP2-O4 (PF3D7_1350900). Expression of these two genes increased approximately 10-fold in artemisinin treated samples (**Table S2**) but not after the other two drug treatments. Additionally, experimental validation such as functional gametocyte measurements or immunofluorescence assays, may also help determine whether the increase in gametocyte formation after artemisinin treatment reflects a true induction of parasite sexual commitment.

Although our results show an enrichment of the expected pathways for atovaquone, we acknowledge some problems with our methodology. For instance, while we aimed to recover 20,000 cells per channel for each 10× Genomics run, the total number of predicted cells obtained was much higher according to Cell Ranger. This suggest a high incidence of multi-species (human and parasite) droplets and thus were filtered for the analyses. Similarly, we obtained a much higher percentage of cells from the first replicate compared to the rest; sequencing mapping showed that those cells were mainly human transcripts rather than *Plasmodium* and thus had to be filtered out. As an attempt to improve this, magnetic enrichment columns were used (second and third replicate) and this did provide an overall higher coverage of the parasite’s genome. This, however, may have introduced a population biased towards later asexual stage rather than obtaining a well-rounded initial population. Additionally, we tried different sequencing depth to evaluate whether it results in more robust results. Indeed, the (third) replicate, with the highest depth and magnetic enrichment, yielded the most complete and high-quality cells. Despite these caveats, all changes highlighted here were consistent and reproducible across different replicates.

Another problem that we faced was the vast difference in parasite population across conditions (*i.e.*, control and drug-treated parasites). We had hypothesized that with single-cell we could bypass the need for synchronization by detecting expression levels in a variety of different stages at once. Indeed, we were able to discern the substages of the asexual lifecycle for all drug-treated parasites, although there were cases where no definitive substage was identified. This could either be related to the drug’s effect, or as an anomaly during cell normalization and integration steps. Based on our results, evaluation of the transcriptomic drug response was only achievable after having a similar number of parasites between the two conditions. Similarly, having to apply two rounds of integration and batch correction steps, while helpful for comparability, could have filtered out rare events or new cell types occurring after treatment specially with clusters having low number of predicted cells. In general, cell filtering and proper cell integration is also imperative for this approach to work.

Despise these difficulties, our results show that single-cell sequencing may serve as a method to investigate the mechanisms of action for uncharacterized molecules, although optimization of the experimental conditions (*e.g.*, initial parasite population, time of drug exposure) must be considered. Additional studies with more compounds/drugs and more drug classes will be needed to further explore this hypothesis.

## Acknowledgments

This work was funded by E.A.W. grants from Gates Foundation (INV-039628) and National Institutes of Health (R01 AI169892, R01 AI172066, R01 AI52533). This publication includes data generated at UC San Diego IGM Genomics Center utilizing an Illumina NovaSeq 6000 that was purchased with funding from a National Institutes of Health SIG grant (#S10 OD026929). The funders had no role in study design, data collection and interpretation, or the decision to submit the work for publication. Authors would like to thank Dyllan Mead, for preparing the parasite cultures.

## Data availability

Raw sequencing generated in this study have been submitted to the NCBI Sequence Read Archive dataset (https://www.ncbi.nlm.nih.gov/sra) under accession number PRJNA1461920. All other data described in this study is available at https://doi.org/10.6084/m9.figshare.32353263 and the **Supplementary Material.**

## Author Contributions

E.A.W. conceived of experiments, reviewed the manuscript, provided funding, performed analyses and designed figures. K.P.G-M. wrote a first draft of the manuscript, performed analyses, constructed supplemental files, and constructed figures. J.C. maintained parasite cultures, performed infections and sequencing. K.J. provided scientific advice and assisted in the experimental design and sequencing. All authors have read and approved the manuscript.

## Materials and Methods

### Plasmodium Dd2 parasite culture and sequencing

*P. falciparum* Dd2 asexual blood-stage parasites were grown in human O-positive (O+) whole blood obtained from the BioIVT and San Diego Blood Bank (SDBB) (BioIVT, HUMANRBCPD-0104976) and was maintained at 2% hematocrit in RPMI 1640 medium (Gibco). The medium was supplemented with 0.25% Albumax II (Gibco), 2 g/liter sodium bicarbonate (Fisher), 0.1 mM hypoxanthine (Sigma), 25 mM HEPES pH 7.4 (Sigma), and 50 μg/liter gentamicin (Gold Biotechnology). To study the drug effect on the parasites, four flasks were generated, one maintained without drug and the remaining three were dosed with one of atovaquone (50nM), GNF179 (100nM), or artemisinin (100nM), respectively. Parasites were incubated at 37°C and 5% CO_2_ in the MIC-101 modular incubator chamber. This process was performed in triplicates. After 48 hours of drug exposure, the culture was passed through a column using magnetic cell separation (MACS, Miltenyi Biotec #130-042-901), where infected red blood cells were retained in the column and washed with non-supplemented RPMI medium (except for replicate 1).

Parasites retained with the magnetic columns were eluted with an RPMI non-supplemented medium, concentrated by centrifugation at 1000 g in a 50μL final volume, and prepared for sequencing following the 10× Genomics™ protocol (Chromium next GEM single-cell 3’ HT Reagent, cat #CG000416). Briefly, 27.7 µL of infected red blood cells (iRBCs; 1,200 cells/µL) were added to the 10× Genomics™ single-cell master mix and loaded onto a Next GEM Chip M aiming to recover 20,000 cells per channel. Single-cell partitioning and barcoding were performed on the Chromium X platform according to the manufacturer’s protocol. Following droplet generation, cDNA within each gel bead-in-emulsion (GEM) was reverse transcribed, amplified, and processed into sequencing ready libraries. Libraries were sequenced on an Illumina NovaSeq 6000 system with S4 flow cells using paired-end reads. Sequencing depth ranged from 100–300 million reads for replicates 1 and 2, and approximately 1.6 billion reads for replicate 3.

### Raw single-cell sequencing data workflow

Raw sequences were manually inspected with FastQC for quality check (https://www.bioinformatics.babraham.ac.uk/projects/fastqc/). A custom reference genome was created with CellRanger(39) *mkref* tool, using the human (GRch38.p13) and *Plasmodium* (PfDd2 v62, PlasmoDB(40)) genomes. Raw sequences were then mapped to the custom genome and transcripts were counted using CellRanger *count* function. Filtered barcoded features were then analyzed using Seurat’s functions(21). Features were first opened with *Read10X* function, and *Plasmodium* features where extracted. From these, protein coding gene transcripts were filtered and used to create a Seurat object.

For each sample (control, artemisinin-, GNF179-, and atovaquone-treated cells), replicate samples were merged and a total of 15,000 cells where randomly selected. Cells where then normalized, highly variable features identified and scaled, and dimensional reduction was performed (PCA) and batch corrected using Harmony(41). To contrast cell conditions, control cells were merged with one of the treated samples (artemisinin, GNF179, and atovaquone). A standard pipeline (highly variable feature detection and scaling, and dimensional reduction transformation) was performed. Cells where then batch corrected and integrated to find the clusters. For each object, cells with less than 50 genes were filtered, and clusters with less than 1,000 cells were removed. Hypervariable gene families (*rifin*, *stevor*, *var*) were filtered from analyses.

### Lifecycle stage cluster identification

For each cluster, we identified the corresponding lifecycle stages using Seurat *FindAllMarkers* function; for each treatment separately, keeping genes with at least 0.25 log_2_ fold-change. We then selected markers having an average log_2_ fold-change of at least 1 in that cluster compared to the rest, expressed in at least 60% of cells in the cluster (pct.1) and no more than 40% in the rest (pct.2), and a *p*-adjusted value of at most 5×10^-5^. In cases where a known stage marker (i.e., an established marker in literature) was present among the list, the corresponding stage was assigned; otherwise, it was left unnamed. All markers were manually verified using the FeaturePlot function in Seurat.

To further detect the lifecycle stages, we compared our markers against the Le Roch *Plasmodium* asexual stage development dataset(3). For this, we selected candidate markers (from *FindAllMarkers*) with at least 30% of cells expressed in the cluster (pct.1) and a *p*-adjusted value of at most 5×10^-5^. For all candidates, the 3D7 orthologs were obtained and matched with the Le Roch (LR) genes. For defining clusters with the matching candidates, we considered LR clusters having more than three matching candidates and assigned the developmental stage based on the most representative known markers or LR established stage (*e.g.,* schizont, gametocyte) for a particular ABS substage. We first focused on LR genes matching to only one cluster in this study (LR gene = *x*, where *x* is the candidate cluster number). For LR genes with “shared” clusters, *i.e.* expressed in two or more candidate clusters, we defined the stage based on the most representative known gene markers across the LR clusters. In the rare cases where we did not obtain a clear stage with the previous criteria, we explored cells expressed at 20% in candidate clusters following the same focus order as described above.

### Differential gene expression analysis

Raw single cell read counts for each sample were extracted using the *AggregateExpression* function from Seurat. Genes with an ortholog in *P. falciparum* 3D7 genome were kept, excluding those with total read counts less than ten. To calculate differential expression, we utilized

DESeq2(42) *DESeqDataSetFromMatrix* with raw counts selecting for one drug treatment versus all control parasites, using the condition (treated, non-treated) as the factor to be tested. This calculation was performed for atovaquone, artemisinin, and GNF179. Normalization and differential calculation were performed using default settings and contrasting by condition. Genes with a *p*-value greater than 5×10^-5^ were filtered.

### Gene Ontology (GO) enrichment analysis

Taking the list of differentially expressed genes unique to each treatment, we split them into two groups: one for upregulated genes (fold-change ≥ 1) and one for downregulated genes (fold-change ≤-1). We filtered genes having a *p-*adjusted value ≤ 0.05 from both lists. Each list was input and queried into PlasmoDB’s gene annotation tool. The “Gene Ontology Enrichment” analysis was performed under the analysis tab, with default settings for biological processes (BP), molecular function (MF) and cellular components (CC). Terms with *P-value* Benjamini corrected < 0.05 were extracted and further analyzed.

