## Supplementary Material for "Single-cell analysis of *Plasmodium falciparum* transcripts after drug perturbation identifies feedback regulation as well as increased transmission potential"

Affiliations
[a]Department of Pediatrics, University of California, San Diego, La Jolla, CA 92093, USA
[b]Institute for Genomic Medicine, University of California, San Diego, La Jolla, CA 92093, USA
[c]Skaggs School of Pharmacy and Pharmaceutical Sciences, University of California, San Diego, La Jolla, CA 92093, USA

[*]Present address: Faculty of Medicine of the National University of San Marcos, Lima, Peru

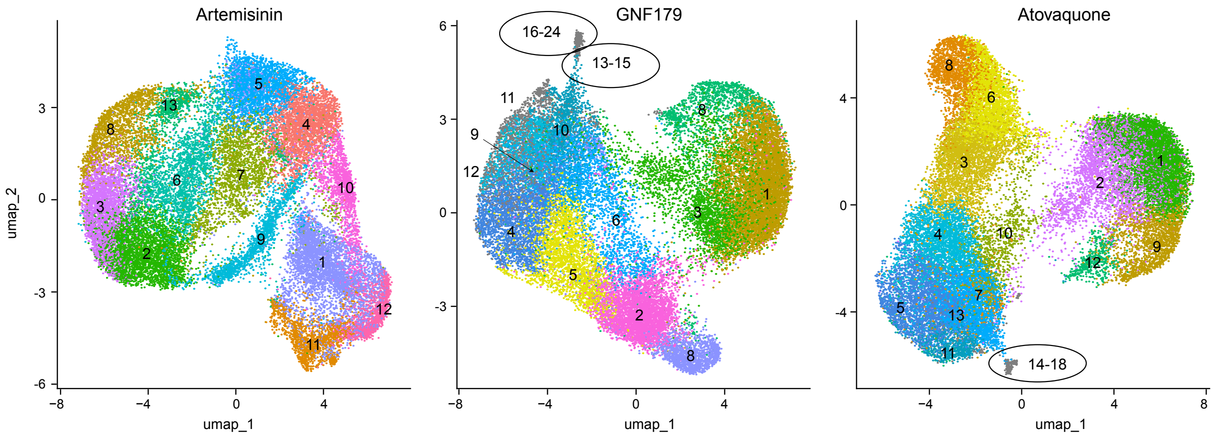

**Supplementary Figure 1. Dimensional reduction clustering for artemisinin, GNF179, and atovaquone treated cells versus non-treated cells.** Plots showing artemisinin, GNF179 and atovaquone dimensional reduction after cell integration and batch correction steps. Clusters were determined using a standard Seurat[1] pipeline (see **Methods**), with resolution of 0.8. Cluster numbers and colors are shown with default settings and represent the total of distinct clusters obtained after using FindNeighbors and FindClusters Seurat's functions. Overlapping clusters (*i.e.*, clusters overlaying between each other) are colored in grey and cluster IDs for three or more overlapping clusters are circled.

| | **Raw set** | | | **Downsampled set** | | |
|---|---|---|---|---|---|---|
| **Condition** | **Mean** | **SD** | **N** | **Mean** | **SD** | **N** |
| Control | 124903.67 | 56212.55 | 3 | 5000 | 2085.27 | 3 |
| Artemisinin | 29174.0 | 29495.06 | 3 | 5001 | 5006.46 | 3 |
| GNF179 | 93705.0 | 66049.06 | 3 | 5000 | 3191.90 | 3 |
| Atovaquone | 91108.33 | 69852.49 | 3 | 5000 | 3147.67 | 3 |

**Supplementary Table 1. Variability between replicates for selected cells across conditions.** Mean number of recovered cells randomly selected across all replicates. Standard deviation is indicated (SD). Values are representative of **Figure 1b**. Calculations performed with GraphPad ("GraphPad Software, Boston, Massachusetts USA, www.graphpad.com").

**Supplementary Table 2. Transcripts mapped to each gene for cells used in the study.**
Attached Excel spreadsheet.

| Cluster | Ctr + artemisinin | | | Ctr + atovaquone | | | Ctr + GNF179 | | | Total |
|---|---|---|---|---|---|---|---|---|---|---|
| | Rep 1 | Rep 2 | Rep 3 | Rep 1 | Rep 2 | Rep 3 | Rep 1 | Rep 2 | Rep 3 | |
| **1** | 227 | 846 | 3400 | 846 | 25 | 3991 | 1416 | 26 | 4819 | 15596 |
| **2** | 522 | 120 | 2803 | 1155 | 155 | 2350 | 418 | 319 | 3400 | 11242 |
| **3** | 505 | 39 | 2269 | 318 | 763 | 2111 | 989 | 180 | 2087 | 9261 |
| **4** | 596 | 1022 | 1085 | 1142 | 1057 | 756 | 2012 | 712 | 207 | 8589 |
| **5** | 764 | 1362 | 572 | 2117 | 665 | 120 | 688 | 991 | 1050 | 8329 |
| **6** | 569 | 204 | 1450 | 261 | 267 | 2354 | 551 | 716 | 1166 | 7538 |
| **7** | 460 | 449 | 1103 | 2025 | 421 | 58 | 347 | 252 | 1160 | 6275 |
| **8** | 908 | 229 | 774 | 312 | 236 | 1605 | 953 | 132 | 357 | 5506 |
| **9** | 581 | 576 | 457 | 989 | 67 | 730 | 1187 | 178 | 23 | 4788 |
| **10** | 6 | 16 | 1498 | 409 | 399 | 303 | 1110 | 226 | 51 | 4018 |
| **11** | 168 | 803 | 507 | 748 | 86 | 7 | 1002 | 158 | 53 | 3532 |
| **12** | 1 | 31 | 1205 | 446 | 52 | 77 | 162 | 421 | 64 | 2459 |
| **13** | 610 | 136 | 454 | 141 | 185 | 15 | 98 | 4 | 0 | 1643 |
| **14** | | | | 95 | 1 | 0 | 41 | 0 | 0 | 137 |
| **15** | | | | 50 | 3 | 0 | 40 | 0 | 0 | 93 |
| **16** | | | | 45 | 0 | 0 | 36 | 0 | 0 | 81 |
| **17** | | | | 22 | 0 | 0 | 34 | 0 | 0 | 56 |
| **18** | | | | 20 | 0 | 0 | 29 | 0 | 0 | 49 |
| **19** | | | | | | | 28 | 0 | 0 | 28 |
| **20** | | | | | | | 24 | 0 | 0 | 24 |
| **21** | | | | | | | 23 | 0 | 0 | 23 |
| **22** | | | | | | | 20 | 1 | 0 | 21 |
| **23** | | | | | | | 20 | 0 | 0 | 20 |
| **24** | | | | | | | 19 | 0 | 0 | 19 |
| **Total** | 5917 | 5833 | 17577 | 11141 | 4382 | 14477 | 11247 | 4316 | 14437 | |

**Supplementary Table 3. Integrated cells across clusters.** Table showing the total number of recovered cells clustered for the downsampled for combined cells with control and artemisinin (29,327), control and atovaquone (30,000), and control and GNF179 (30,000). Total number of cells mapping to each cluster per replicate were determined with Seurat and split based on replicate number. The set of integrated cells with artemisinin and control parasites had an artifact during the integration resulting in lower cell count. Clusters with cells predominately from one replicate compared to the rest were filtered, namely clusters 14–18 for atovaquone (replicate 1) and 9–11 and 13–24 from GNF179-treated cells (replicate 1). Clusters with less than 1,000 cells with noticeable differences between replicates were filtered, including 11–13 from atovaquone-treated cells and substage 12 from GNF179-treated cells.

**Supplementary Table 4. Asexual life cycle stage identification.** Attached Excel spreadsheet.

| Treatment | Cluster | Gene | $Log_2$ fold change | P-value |
|---|---|---|---|---|
| Artemisinin | Ring or Trophozoite 1 vs 2 | *pfmesa* (mature parasite-infected erythrocyte surface antigen); adjacent to trophozoite | 2.02 | $1.00x10^{-21}$ |
| | | *pfsera-5* (serine repeat antigen 7); adjacent to schizont | -2.40 | $7.72x10^{-165}$ |
| | Schizont 1 vs 2 | *pfH-3* (histone H3); adjacent to ring/trophozoite | 1.18 | $2.70x10^{-46}$ |
| | | *pfetramp-11.2* (early transcribed membrane protein 11.2); adjacent to ring | -2.36 | $8.58x10^{-153}$ |
| GNF179 | Ring or Trophozoite 1 vs 2 | *pfrhopH-2* (high molecular weight rhoptry protein 2); adjacent to schizont | 1.94 | $2.38x10^{-127}$ |
| Atovaquone | Ring 1 vs 2 | *pfepf-1* (exported protein family 1); adjacent to trophozoites | 4.39 | $6.05x10^{-188}$ |
| | | *pfgarp* (glutamic acid-rich protein); adjacent to schizont/merozoite | -2.92 | $4.52x10^{-121}$ |
| | Schizont or Merozoite 1 vs 2 | *pfama-1* (apical membrane antigen 1); adjacent to schizont | 1.20 | $1.02x10^{-97}$ |

**Supplementary Table 5. Fold-change differences between same stage clusters for GNF179, atovaquone, and artemisinin treatments.** Gene expression was calculated with Seurat's *FindMarkers*. Clusters were compared according to color code in **Figure 2a**. Cluster 1 was set as *x* and the remaining as cluster(s) as *y*, in the *FindMarkers* function. *P*-values were calculated according to *t*-test and Bonferroni corrected. The most representative known marker from each cluster was selected to highlight the neighboring parasite stage clusters, except for atovaquone ring three versus the rest, where it was selected based on Le Roch et al. expression data[2].

**Supplementary Table 6. Estimated number of unique single-cell cells per cluster parasite stage before and after drug-treatment.** Attached Excel spreadsheet.

**Supplementary Table 7. Differentially expressed genes for drug-treated versus control cells.**
Attached Excel spreadsheet.

**Supplementary Table 8. Gene ontology analysis for differentially expressed genes after GNF179, atovaquone, and artemisinin treatment.** Attached Excel spreadsheet.
